# Stable representation of sounds in the posterior striatum during flexible auditory decisions

**DOI:** 10.1101/180505

**Authors:** Lan Guo, William I. Walker, Nicholas D. Ponvert, Phoebe L. Penix, Santiago Jaramillo

**Author notes:** Co-first author (Lan Guo, William I. Walker).

## Abstract

The neuronal pathways that link sounds to rewarded actions remain elusive. It is unclear whether neurons in the posterior tail of the dorsal striatum (which receive direct input from the auditory system) mediate action selection, as other striatal circuits do. Here, we examined the role of posterior striatal neurons in auditory decisions in mice. We found that, in contrast to the anterior dorsal striatum, activation of the posterior striatum did not elicit systematic movement. However, activation of posterior striatal neurons during sound presentation in an auditory discrimination task biased the animals’ choices, and transient inactivation of these neurons largely impaired sound discrimination. Moreover, the activity of these neurons reliably encoded stimulus features, but was only minimally influenced by the animals’ choices. Our results suggest that posterior striatal neurons play an essential role in auditory decisions, yet these neurons provide sensory information downstream rather than motor commands during well-learned sound-driven tasks.

## Introduction

In the mammalian brain, the dorsal striatum links neural signals from the cerebral cortex to circuits in the basal ganglia to mediate action selection. Electrophysiological and inactivation studies have identified two regions within the dorsal striatum which play distinct roles in decision making: the dorsomedial striatum (DMS) involved in flexible goal-oriented behavior, and the dorsolateral striatum (DLS) which mediates habitual actions (Yin and Knowlton, 2006; Balleine et al., 2009; Devan et al., 2011). Recent anatomical characterization of the excitatory input from cortex and thalamus onto the striatum suggests that the organization of the dorsal striatum goes beyond the DMS and DLS divide (Hunnicutt et al., 2016). This characterization in rodents showed that the posterior portion of the striatum receives a combination of sensory inputs that sets it apart from other regions. Similarly, an evaluation of reward-related signals of the dopaminergic input along the anterior-posterior axis of the striatum provides further evidence that the posterior tail of the striatum forms a circuit distinct from DMS and DLS (Menegas et al., 2017). It is not clear, however, whether the function of this posterior region is qualitatively different from the previously characterized striatal subregions. Here, we evaluate the role of neurons in the posterior tail of the striatum during sensory-driven decisions in mice.

The posterior tail of the dorsal striatum in rodents receives direct neuronal projections from the auditory thalamus (ATh) and the auditory cortex (AC), as well as midbrain dopaminergic signals (Menegas et al., 2015; Hunnicutt et al., 2016). Because of these anatomical features, this region is sometimes referred to as the auditory striatum (Znamenskiy and Zador, 2013). Given this convergence of sensory and reward-related signals, and prompted by the role of other dorsal striatal regions, we hypothesized that the tail of the striatum drives rewarded actions according to acoustic cues. Here we show that such a hypothesis does not fully account for the role of this striatal region during sound-driven decisions. In summary, we found that posterior striatal neurons were necessary for the expression of sound-action associations, and that stimulation of these neurons biased decisions based on sounds. In contrast to activation of anterior dorsal striatal neurons, activation of posterior striatal neurons did not drive movement. Moreover, when a behavioral task required rapid updating of sound-action associations without changes in the expected reward, the representation of sounds by the large majority of posterior striatal neurons was stable across contexts and did not depend on the animal’s choice. These results suggest that in a well-learned sound-driven decision task, neurons in the posterior striatum provide sensory information downstream, while providing little information about behavioral choice before action initiation.

## Results

### Activation of distinct subregions of the dorsal striatum produces different effects on movement

The striatum is comprised of two main neuronal outputs, the direct (or striatonigral) pathway and the indirect (or striatopallidal) pathway. One experimentally-supported model of dorsal striatal function posits that the striatal direct pathway drives action initiation (Kravitz; et al., 2010; Freeze et al., 2013). To test whether activation of the posterior striatum produces similar effects on motor initiation as the anterior (dorsomedial) striatum, we used Drd1a::ChR2 mice which express channelrhodopsin-2 (ChR2) in direct pathway medium spiny neurons (dMSNs), and optogenetically activated these neurons in freely-moving animals (Figure 1A).

**Figure 1.**
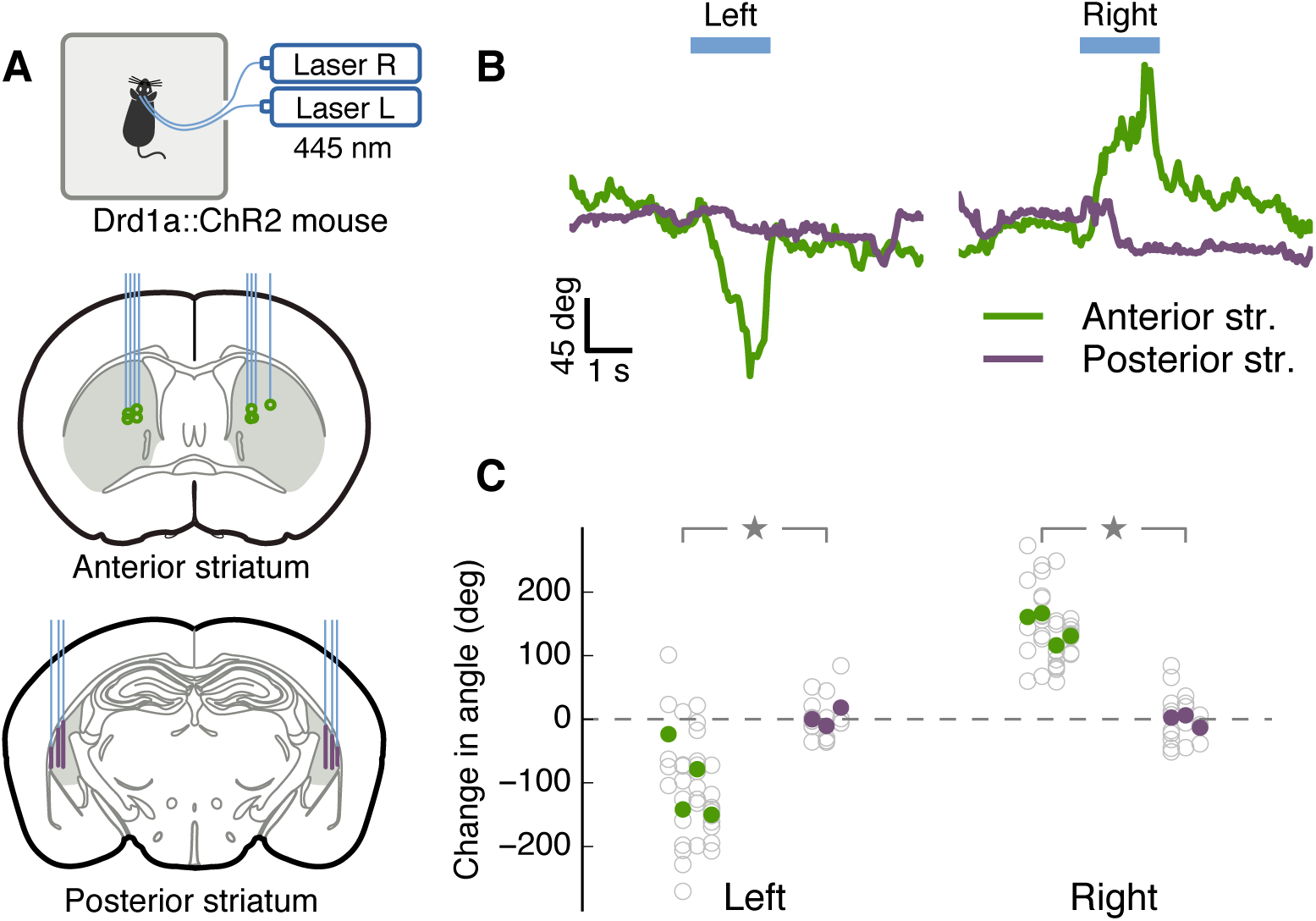
Activation of distinct subregions of the dorsal striatum produced different effects on movement. (A) Top: experimental setup. Optogenetic stimulation in freely moving mice of direct-pathway neurons from one of four different sites in the dorsal striatum: anterior striatum (left or right) and posterior striatum (left or right). Middle: Coronal brain slice. Green dots indicate the tip of fixed optical fibers implanted in the anterior striatum (gray) confirmed postmortem. Bottom: purple lines indicate the stimulation sites by movable optical fibers implanted in the posterior striatum. (B) Representative head angle trace over one trial of unilateral stimulation at each site. The blue bar represents the laser pulse (1.5 seconds) delivered in each trial. Positive angles correspond to left rotation. (C) Average change in head angle by optogenetic stimulation in each mouse tested. Each gray circle is one trial, each filled circle is the average for one hemisphere of one mouse. Stimulation of anterior striatum (green) produced a significantly larger change in head angle compared to stimulation of the posterior striatum (purple) in either hemisphere (*p* = 0.034, Wilcoxon rank-sum test).

We found that unilateral optogenetic stimulation of anterior dorsal striatal dMSNs in freely-moving mice elicited contralateral head rotation clearly visible in single trials (Figure 1B, green trace). In contrast, unilateral stimulation of the same magnitude and duration in the posterior striatum did not result in head or body rotation (Figure 1B, purple trace). Analysis of each animal showed a clear difference between stimulating the left *vs.* right hemisphere in the anterior striatum (*p <* 0.05 for each of 4 mice, Wilcoxon rank-sum test), but no significant effect in the posterior striatum (*p >* 0.1 for each of 3 mice, Wilcoxon rank-sum test). Across the mice tested, the average head rotation to the contralateral side of stimulation was 121*±*45 degrees (mean*±*SD) for dorsomedial striatum and 4*±*10 degrees for posterior striatum. This difference was statistically significant on each hemisphere (Figure 1C, left hemi: *p* = 0.034; right hemi: *p* = 0.034, Wilcoxon rank-sum test).

These results indicate that activation of direct pathway neurons in the posterior striatum does not directly drive movement, in contrast to other dorsal striatal regions. Because the posterior striatal neurons receive dense auditory input, we set out to evaluate the function of these neurons in the context of sound-driven decisions.

### Activation of direct-pathway posterior striatal neurons biases sound-driven decisions

To test the role of posterior striatal neurons in sound-driven decisions, we first evaluated the effects of activating posterior striatal dMSNs during a two-alternative choice sound discrimination task. Mice initiated a trial by poking their nose in the center port of a 3-port chamber. After the presentation of a 100 ms sound, mice were required to go to either the left or right reward port based on the frequency of the acoustic stimulus (Figure 2A). To test the effect of activating posterior striatal dMSNs in the task, the same group of mice in which optogenetic stimulation did not elicit rotational movement (Figure 1) were tested while they performed the task. Optogenetic stimulation for 200 ms (50 msec before sound onset to 50 msec after sound offset) was presented in 20% of the trials. Unilateral activation of posterior striatal dMSNs during sound presentation resulted in a contralateral choice bias apparent in single sessions (Figure 2B,C). Average performance was clearly different between trials with stimulation of the left hemisphere and stimulation of the right hemisphere (*p <* 0.001, Wilcoxon rank sum test, Figure 2D). This effect was significant for each Drd1a::ChR2 mouse tested (*p <* 0.01 for each of 3 mice, Wilcoxon rank-sum test). No effect was observed when the same procedure was applied to wild-type mice, confirming that the observed bias was not a result of visible light stimulation (Figure S1).

**Figure 2.**
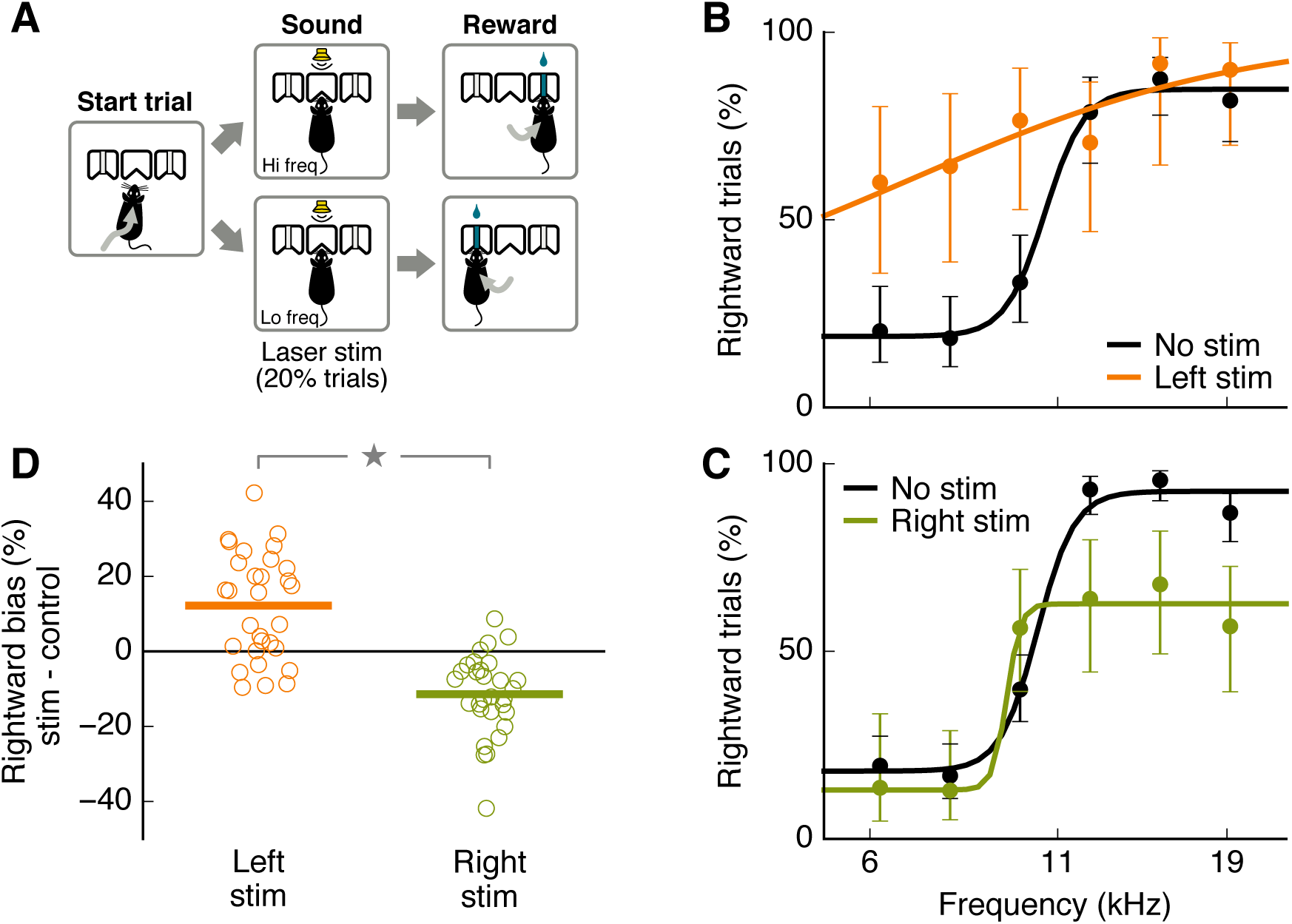
Activation of direct-pathway posterior striatal neurons biased sound-driven decisions. Schematic of the two-alternative choice sound frequency discrimination task. Mice initiated each trial by entering a center port and had to choose one of two side reward ports depending on the sound presented: low frequency = left, high-frequency = right. (B) Psychometric performance for one behavioral session that included optogenetic activation of direct-pathway neurons in the left posterior striatum on 20% of trials. Error bars indicate 95% confidence intervals. (C) Psychometric performance for one behavioral session that included optogenetic activation of direct-pathway neurons in the right posterior striatum. (D) Change in the percentage of rightward choices during optogenetic stimulation for each hemisphere. Each dot represents one session (N=3 mice, 10 sessions each hemisphere per mouse). Horizontal bars represent averages across all sessions for all mice. Stimulation produced significantly different biases in the left versus the right hemisphere (*p <* 0.001, Wilcoxon rank-sum test).

To test whether the observed bias depended on the stimulated site within a hemisphere, we first examined the magnitude of stimulation effects based on fiber location along the medial-lateral axis of the posterior striatum. We found that activation of regions in the border between striatum and cortex resulted in significantly lower bias than activation in the center of the striatum (Figure S1). In addition, we tested whether the bias direction depended on the frequency tuning of the stimulated site. We measured the sound response of each stimulated site through a tetrode bundle implanted alongside the optical fiber. Both hemispheres contained sites responsive to sound frequencies above or below the sound categorization boundary. We found no significant correlation between the preferred frequency of a stimulated site and the direction of the observed behavioral bias (*r* = 0.135, *p* = 0.478, Spearman correlation test, Figure S2). A contralateral bias without clear frequency-specific bias is reminiscent of the effects observed when stimulating the auditory cortex without cell-type specificity (supplementary material of Znamenskiy and Zador (2013)).

These results indicate that activation of posterior striatal neurons during the sound presentation biases decisions in the auditory task, even though similar stimulation does not drive movement outside the task. We next set out to test the necessity of these neurons for sound-driven decisions.

### Inactivation of posterior striatal neurons impairs sound-driven decisions

To test whether the activity of posterior striatal neurons was required when performing the sound-discrimination task, we quantified task performance during bilateral reversible inactivation of these neurons (Figure 3, Figure S3). Injection of muscimol (a GABA-A receptor agonist) in the posterior striatum resulted in a consistent decrease in task performance compared to injection of saline as control (Figure 3B). The effect was observed in all mice tested (Figure 3C, *p <* 0.05 for each mouse, Wilcoxon rank-sum test), and consisted of a flattening of the psychometric curve with minimal impairments on the animal’s ability to initiate trials or collect reward. These results were replicated using fluorescent muscimol to confirm that inactivation restricted to the posterior striatum affected task performance (Figure S4). Even though animals performed fewer trials during muscimol inactivation sessions compared to saline control sessions (*p <* 0.05 for three out of five mice, Wilcoxon rank-sum test), they still performed hundreds of trials during each session (480*±*146 muscimol *vs.* 733*±*86 saline, Figure S5). As a result of muscimol inactivation, mice displayed slower withdrawals from the center port on average (*p <* 0.05 for four out of five mice, Wilcoxon rank-sum test, Figure S5). A change in movement speed from the center port to the reward ports was observed in only one out of five mice (*p <* 0.05, Wilcoxon rank-sum test, Figure S5). During inactivation sessions, some animals used a strategy in which they chose a reward port at random, while other animals displayed a strong bias to one side. Both of these strategies resulted in an average performance close to chance level for the binary choice.

**Figure 3.**
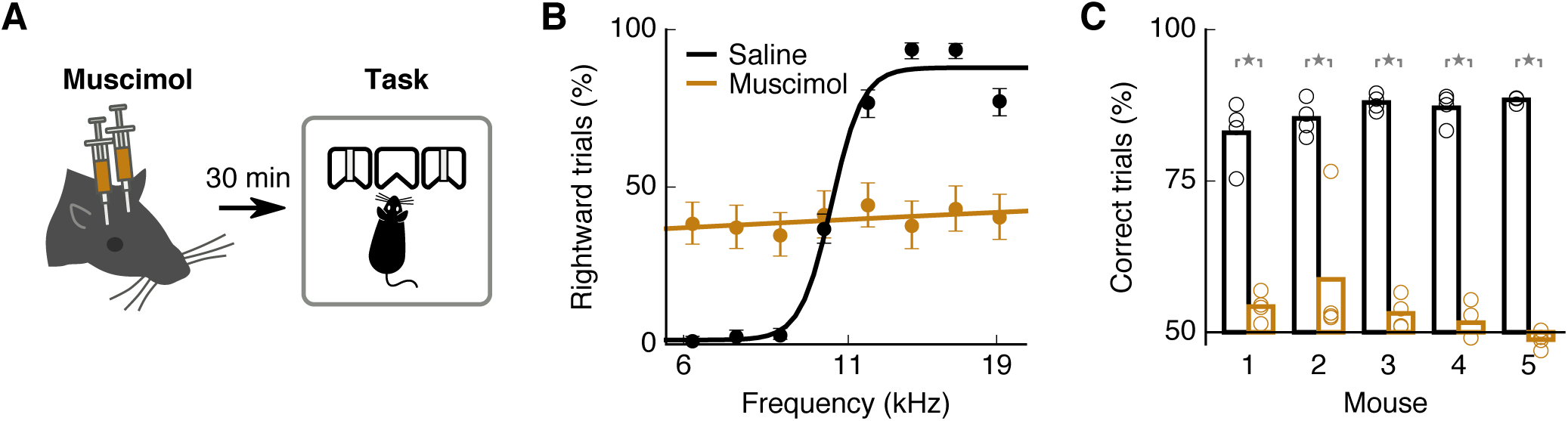
Inactivation of posterior striatal neurons impaired sound-driven decisions. (A) Mice were injected bilaterally with muscimol in the posterior striatum 30 minutes before they performed the two-alternative choice sound discrimination task. (B) Average psychometric performance for one mouse on sessions with injection of muscimol (4 sessions) or saline control (4 sessions). Error bars indicate 95% confidence intervals. (C) Average percentage of correct trials on each saline session (black) and each muscimol session (brown) for each mouse. Bars indicate average across sessions for each mouse. Muscimol inactivation significantly reduced the percentage of correct trials on each mouse (*p* = 0.021, Wilcoxon rank-sum test).

These inactivation results indicate that the activity of neurons in the posterior striatum is necessary for auditory decisions, but not for executing the movements required by the task. We next quantified what information is encoded by posterior striatal neurons during sound-driven decisions.

### Activity of posterior striatal neurons correlates with sounds and actions

To further delineate the possible functional roles of posterior striatal neurons, we recorded activity from single neurons and examined their response to different components of the sound-driven decision task. We first evaluated the neuronal responses to sound stimuli presented in the two-alternative choice task.

We found clear neural responses to sounds and selectivity to sound frequency in neurons from the posterior striatum. Figure 4A,B shows sound responses from two distinct neurons during the discrimination task. To estimate how many neurons were responsive to sounds during the task, we compared the firing rate of each neuron during the sound presentation (all trials pooled together) against the neurons spontaneous firing (Figure 4C). We found that 44.6% (232/520) of cells showed a significant change in firing in response to sound (*p <* 0.05, Wilcoxon rank-sum test). We then quantified how many neurons provided sound information relevant to the task by comparing sound evoked responses between high frequency (any sound above the categorization boundary) *vs.* low frequency trials (any sound below the boundary). We found that 26.3% (137/520) of all cells showed a different average response between high- and low-frequency sounds during the task (*p <* 0.05, Wilcoxon rank-sum test, Figure 4D). We further characterize the frequency selectivity of neurons in the posterior striatum by comparing the activity evoked by each sound presented in the task. From all recorded cells, 38.8% (202/520) responded differently across stimuli (*p <* 0.05, one-way ANOVA). Sound responses and frequency selectivity were also observed when sounds were presented outside the context of the discrimination task (Figure S6). These measurements indicate that the identity of sounds has a strong influence on the firing rate of neurons in the posterior striatum.

**Figure 4.**
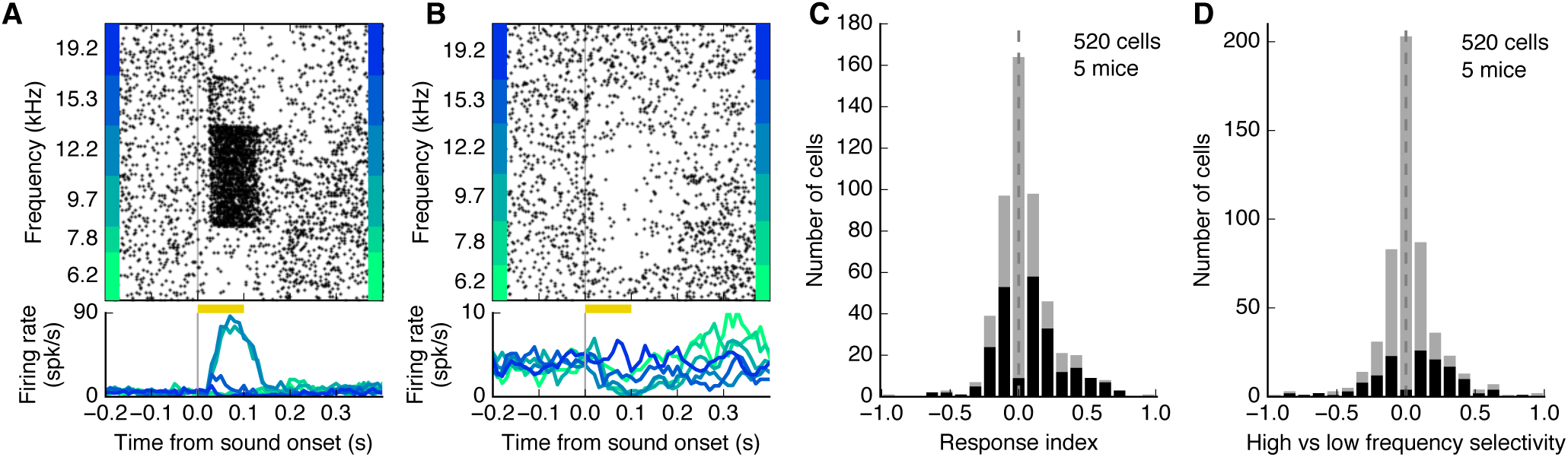
Posterior striatal neurons displayed frequency-selective sound-evoked responses. (A) Example of sound responses from a posterior striatal neuron during the sound discrimination task. Yellow bar indicates the duration of the sound. (B) Sound responses of a different posterior striatal neuron which showed only suppression of activity. Both example cells showed clear frequency selectivity. (C) Magnitude of sound-evoked response for each neuron (all trials pooled together). A positive response index indicates an increase in activity compared to baseline spontaneous activity. A negative index, a decrease in activity. Neurons with statistically significant evoked responses (*p <* 0.05, Wilcoxon rank-sum test) are shown in black. (D) Sound selectivity index calculated by comparing neural responses to high *vs.* low frequency sounds based on the categorization boundary from the task. Neurons with statistically significant differences (*p <* 0.05, Wilcoxon rank-sum test) are shown in black.

To examine whether the activity of posterior striatal neurons was correlated with the movement of the animals, we quantified the difference in firing rate during movement toward the left port *vs.* the right port. After an animal left the center port, that activity of neurons was often different depending on movement direction (Figure 5A,B). We found that this effect was significant (*p <* 0.05, Wilcoxon rank-sum test) for 38.5% (200/520) of all cells recorded (Figure 5C). Of the subset of cells that were selective to movement direction, 55.5% showed stronger firing during movement contralateral to the recording hemisphere. Across the whole population, more cells fired at higher rates during movement contralateral to the recording site (*p* = 0.025, Wilcoxon signedrank test). Last, we evaluated the relation between left/right movement selectivity and low/high sound frequency selectivity. We found that 11.9% (62/520) of neurons were both selective to sound frequency and movement direction during the behavioral task (Figure S7).

**Figure 5.**
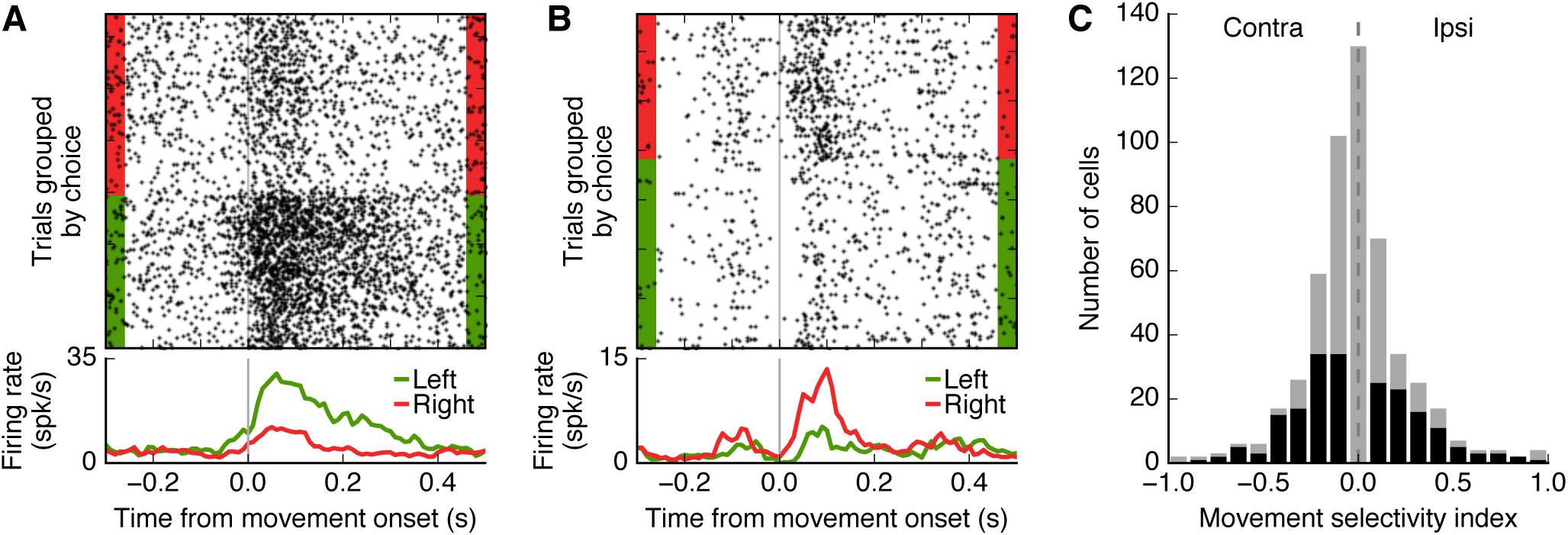
Firing rate of posterior striatal neurons during movement depends on movement direction. (A) Activity from a posterior striatal neurons aligned to the moment the mouse left the center port and moved toward a reward port. Activity differed between right and left choices. Plot includes all trials (any stimulus frequency). (B) Activity from a different posterior striatal neuron showing the opposite movement selectivity. (C) Movement selectivity index: (I-C)/(I+C), where I and C are the average firing rates in the period 50-150 ms after leaving the center port for choices ipsilateral and contralateral to the recording hemisphere, respectively. N = 520 cells recorded from 5 mice. Cells with a statistically significant difference in activity during contralateral and ipsilateral choices (*p <* 0.05, Wilcoxon rank-sum test) are shown in black.

These results indicate that at different time periods during the behavioral task, posterior striatal neurons encode information about sounds and actions. However, these observations are not sufficient to know whether these neurons encoded the sensory-motor association during the sound presentation, or simply relay auditory information before movement initiation. We therefore tested whether the neuronal activity during sound presentation was influenced by the animal’s choice.

### Behavioral choice influences sound-evoked activity in a subset of posterior striatal neurons

One possible model for auditory decision-making assumes that the posterior striatum is a main locus of action selection, given sensory information from cortex and thalamus. This model predicts that sound-evoked responses of a striatal neuron will differ depending on the animal’s action, even if the stimulus is the same. To test this prediction, we compared neural activity evoked by the same sound between trials in which a mouse chose the left reward port and trials in which the mouse chose the right reward port.

We focused our analysis on trials with sound stimuli near the categorization boundary for which animals performed near 50% accuracy, and the number of left and right choices was about the same. Figure 6A shows a posterior striatal neuron in which sound-evoked responses were influenced by the animal’s choice. However, for most neurons, the animal’s choice had little to no influence on the evoked-responses, as in the example neuron shown in Figure 6B. From our analysis of posterior striatal neurons that responded to stimuli near the categorization boundary, we found that 11.5% (7/61) had a significant change in evoked response depending on the animal’s choice (Figure 6C, *p <* 0.05, Wilcoxon rank sum test). We found no systematic relation between the spike shape of each neuron and the magnitude of its response modulation by choice (Figure S8). Moreover, the magnitude of modulation by choice was not correlated to the magnitude of movement selectivity in these cells (Figure S9, *r* = 0.02, *p* = 0.8, Spearman correlation test).

**Figure 6.**
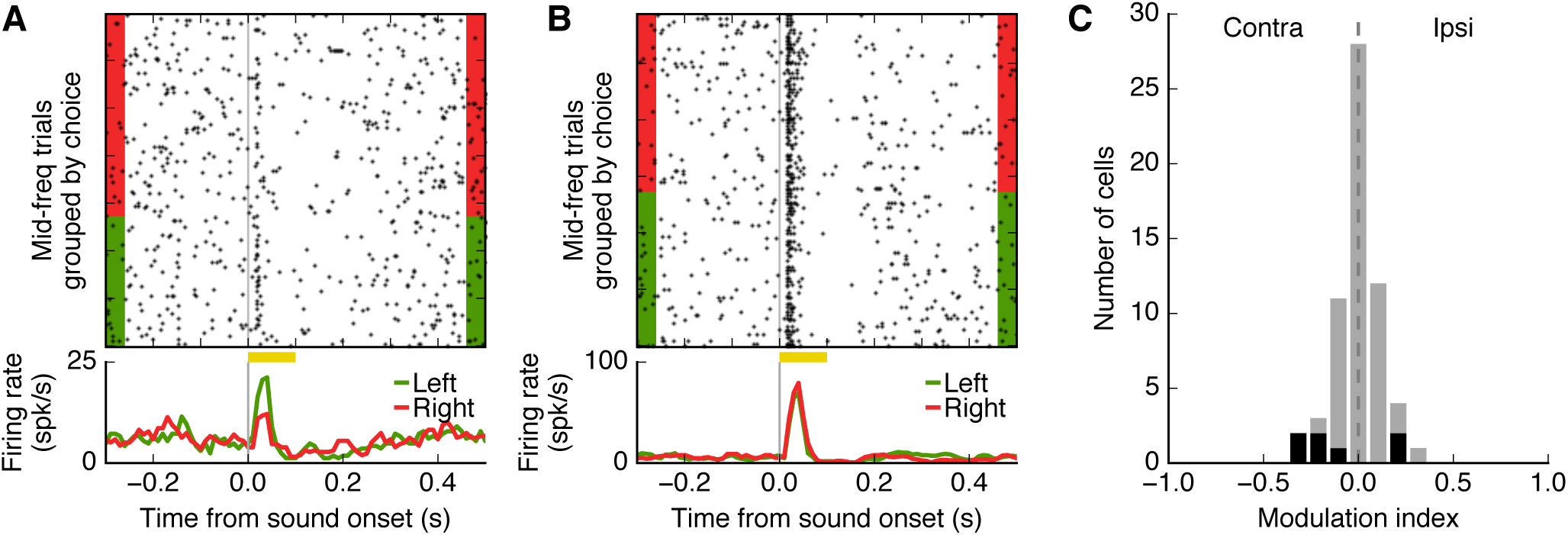
Choice influenced sound-evoked responses in a subset of posterior striatal neurons. (A) Response of one neuron to a stimulus near the categorization boundary, grouped according to the animal’s choice. Sound-evoked response for this neuron was modulated by the animal’s choice. Activity of a different neuron showing no influence of choice on the sound-evoked response. Influence of choice on sound-evoked activity for all neurons that showed a response to stimuli near the categorization boundary (N = 67 cells from 5 mice). Less than 12% of neurons showed a significant modulation by choice (*p <* 0.05, Wilcoxon rank-sum test), shown in black.

To test whether posterior striatal neurons encoded the animal’s choice while in the center port, we also quantified the activity of each neuron during the 100 ms window before mice left the center port. This analysis included only trials with the stimulus near the boundary, to control for the changes in activity due to sound frequency. We found that 4.6% (24/520) of neurons displayed significantly different activity before action initiation based on the subsequent choice (Figure S10, *p <* 0.05, Wilcoxon rank sum test).

These results suggest that in the case of ambiguous (difficult) stimuli, activity of posterior striatal neurons before action initiation contains little information about the animal’s subsequent choice. However, these results do not rule out the possibility that information about choice would be present in sound-evoked responses when an animal has formed a clear association between a stimulus and an action.

### Effects of rapid changes in sound-action associations on activity of posterior striatal neurons

In the experiment described in the previous section, the animals’ variability in choice arose from the difficulty of perceptual decisions near the decision boundary. We hypothesized that a larger number of neurons would be influenced by the animal’s choice if instead of using ambiguous stimuli, the meaning of the stimuli changed systematically from predicting reward on the left port to predicting reward on the right port.

To test this hypothesis, we used a previously developed task for rodents (Jaramillo and Zador, 2014) in which the rewarded action associated with a subset of sounds changes every few hundred trials (Figure 7A). In this switching task, low-frequency sounds still indicate reward on the left side (and high-frequency indicates reward on the right), but the categorization boundary changes across blocks of trials. As a result, a stimulus of intermediate frequency (*e.g.*, 11 kHz) is associated with the left port in one block of trials and associated with the right port on the next block. Importantly, the amount of reward associated with such sound remains the same, and it is only the action associated with this sound that changes. Mice trained in this flexible categorization task achieved high performance levels and were able to switch between sound-action association contingencies several times per session, as previously reported (Jaramillo and Zador, 2014).

**Figure 7.**
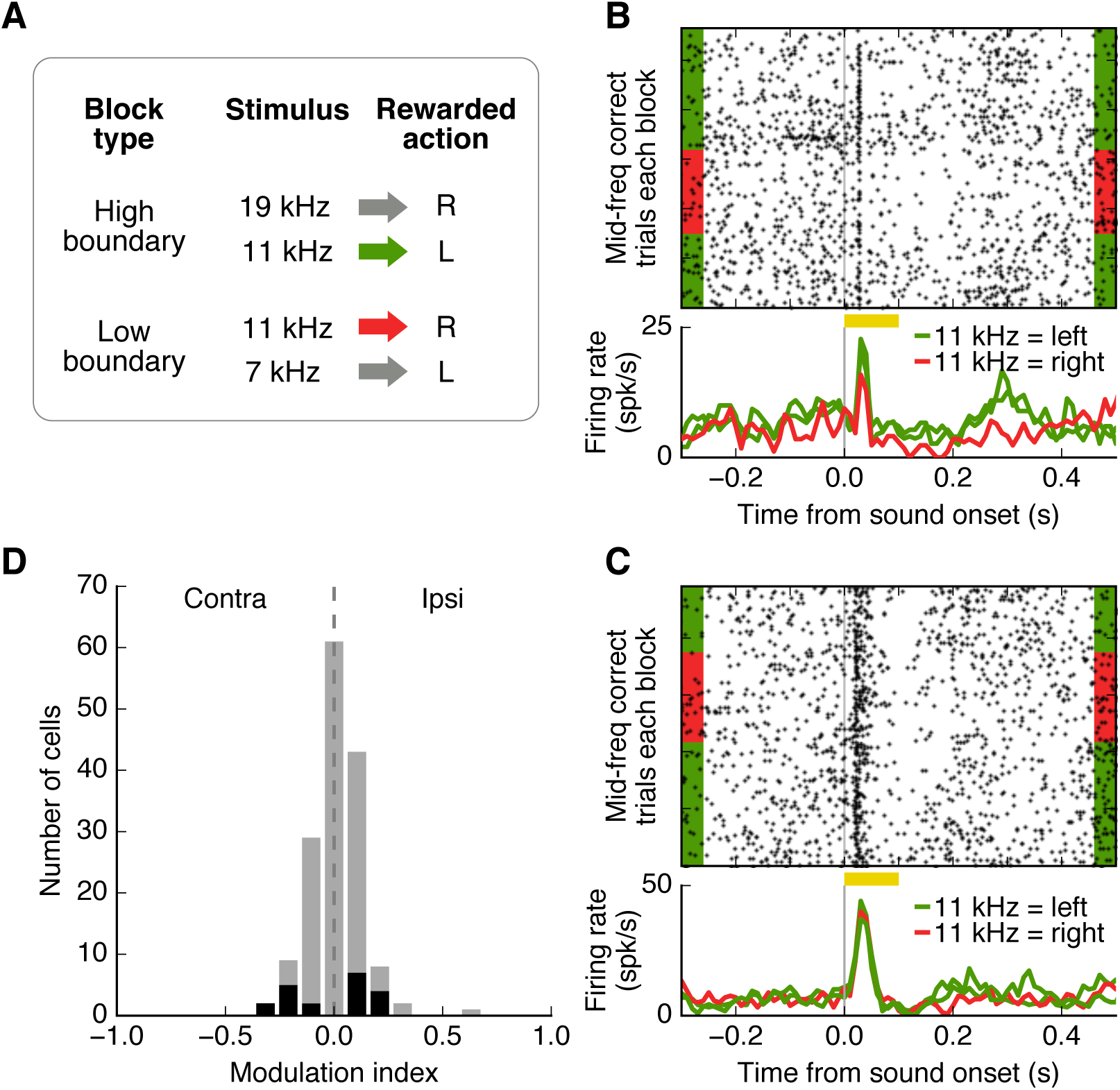
Effect of rapid changes in sound-action associations on activity of posterior striatal neurons. (A) Schematic of the switching task. The rewarded action associated with a sound of intermediate frequency changed from one block of trials to the next. (B) Responses of one posterior striatal neuron to the stimulus of intermediate frequency for three blocks of trials. Only correct trials are included. Sound-evoked responses for this neuron changed systematically depending on the rewarded action associated with the stimulus. (C) Activity of a different neuron showing no change in sound-evoked responses across sound-action contingencies. (D) Influence of changing sound-action association on sound-evoked activity for all neurons that showed a response to the switching stimulus (N = 155 cells from 4 mice). Less than 13% of neurons showed a significant change across sound-action contingencies (*p <* 0.05, Wilcoxon rank-sum test), shown in black.

We found that, in a subset of posterior striatal cells, the evoked-response to a given sound changed depending on the port associated with that sound (Figure 7B). However, most cells showed a stable representation of sounds across blocks of trials, independent of the associated reward port (Figure 7C). From our analysis of neurons that responded to the stimulus of intermediate frequency, we found that sound-evoked responses in 12.9% (20/155) of cells were influenced by changes in the sound-action association (Figure 7D, *p <* 0.05, Wilcoxon rank-sum test). Similarly, when we evaluated neuronal activity before mice left the center port in these trials, 7% (51/725) of cells showed a statistically significant change in activity according to the animal’s choice (Figure S10).

These results provide further support for the idea that posterior striatal neurons display a stable representation of sounds, minimally affected by action selection. Under the conditions tested, the activity of posterior striatal neurons encoded information about sound identity, rather than the behavioral choice or the sound-action association.

## Discussion

The main objective of this study was to evaluate the role of neurons in the posterior tail of the rodent striatum during sensory-driven decisions. Our experiments helped evaluate which roles these neurons play in the pathway linking sensation to action: from representing the raw stimulus, to performing computations necessary for stimulus discrimination, choice selection or action execution. Based on previous studies of the dorsal striatum (Kravitz; et al., 2010; Freeze et al., 2013; Sippy et al., 2015), together with evidence of synaptic plasticity in the cortico-striatal pathway during reward-driven learning (Reynolds et al., 2001; Xiong et al., 2015), we posited that posterior striatal neurons would drive actions according to rewarded associations to acoustic stimuli. We found, however, that after animals have learned a sound discrimination task, the activity of these neurons before action initiation contained more information about the identity of the stimulus than about the animal’s choice.

### A sensory subregion of the dorsal striatum

In contrast to the fine parcellation commonly used to describe the cerebral cortex, a functional sub-division of the dorsal striatum is usually limited to a lateral region (involved in habitual behaviors) and a medial region (involved in flexible behaviors). However, studies in mouse and rat evaluating the inputs onto striatal circuits from cortical, thalamic and dopaminergic neurons demonstrate a more refined anatomical organization of the dorsal striatum which in turn suggests the existence of further functional specialization (McGeorge and Faull, 1989; Glynn and Ahmad, 2002; Hunnicutt et al., 2016; Menegas et al., 2017). A characterization of the distribution of dopamine receptors across the rostro-caudal axis provides further support for this refinement (Gangarossa et al., 2013). In particular, the posterior portion of the rodent striatum has been identified as an area that receives convergent inputs from sensory cortex, sensory thalamus, and midbrain dopamine neurons.

Our results in mice demonstrate that a large fraction of neurons in this region of the striatum have clear selectivity to sound features, consistent with previous observations in the rat (Bordi and LeDoux, 1992; Znamenskiy and Zador, 2013). In our sound discrimination task, over a quarter of the recorded posterior striatal neurons were able to distinguish between sound frequencies above and below the categorization boundary, and a larger percentage discriminated across different stimuli (even within a category). These finds suggest that the fine sound frequency selectivity in posterior striatal neurons does not result simply from associating a sound to left *vs.* right actions. In addition, when tested with stimuli close to the boundary, we found little information about the animals’ choices encoded in the neural activity during the presentation of the sound. Although our task does not allow for an accurate estimate of the moment the decision is made, our analysis of activity before movement initiation also yielded a minimal influence by choice. We conclude therefore that very few neurons in this brain region (less than 13% in this study) encode the sound-action association on each trial. This result is comparable to observations from the auditory thalamus and auditory cortex of rats during a similar task (Jaramillo et al., 2014), which raises the possibility that activity changes observed in the striatum simply reflect modulated signals arriving from thalamic or cortical inputs. Taken together, these results support a model in which posterior striatal neurons provide sensory information rather than encoding the behavioral choice in well-learned tasks.

### The sensory striatum as an essential pathway for auditory decisions

Lesion studies indicate that rodents can perform sound-action association tasks without the auditory cortex (Teich et al., 1989; Campeau and Davis, 1995; Song et al., 2010; Gimenez et al., 2015). In contrast, lesions of the auditory thalamus largely affect expression of fear conditioning to sounds (Campeau and Davis, 1995) and discrimination of sound frequencies (Gimenez et al., 2015). These observations suggest that outputs of the auditory thalamus convey signals essential for auditory decisions. The dorsal striatum, a direct target of the auditory thalamus with connections to motor structures, is therefore a likely candidate circuit to mediate tasks that require sound-driven decisions.

Our inactivation experiments demonstrate that silencing auditory striatal neurons has a drastic effect on a task that requires sounds-driven decisions. One interpretation of these results states that these neurons belong to a unique pathway required for successful performance of the frequency discrimination task we study. Alternatively, these neurons could form one of several parallel pathways linking sensation to action, and reversible inactivation yields strong effects on behavior by altering the dynamics of downstream circuits (Otchy et al., 2015). Our experiments cannot distinguish between these possibilities, yet, provide strong support for a role of these striatal neurons in sensory-driven decisions.

### The role of striatal neurons in auditory decisions

In contrast to the results of activating the anterior dorsal striatum, optogenetic activation of direct-pathway neurons in the posterior striatum did not elicit movement outside the behavior task, indicating that these cells play a distinct role in motor initiation from neurons in the anterior portion. Based on these results and the sound selectivity we observed, we postulated that activation of posterior striatal dMSNs would bias animals according to the preferred frequency of the stimulated cells. In fact, activation of rat auditory cortico-striatal neurons produced a choice bias that depended on the frequency tuning of the stimulated site (Znamenskiy and Zador, 2013). In the same study, however, optogenetic stimulation of auditory cortical neurons without cell-type specificity caused a contralateral choice bias that did not depend on frequency tuning (see supplementary figure S3 of Znamenskiy and Zador (2013)). Our results when stimulating dMSNs in the posterior striatum recapitulated the latter finding by producing a consistent contralateral bias with only a weak (and not statistically significant) correlation to stimulation site tuning. Given that both direct- and indirect-pathway MSNs receive input from cortico-striatal neurons, our results do not rule out the possibility that a balance between dMSN and iMSN activity in the posterior striatum is required to produce a frequency-specific bias.

All measurements in our study were performed after animals had achieved high performance in the discrimination task. Therefore, our data does not provide evidence for the role of the posterior striatal circuits during learning. A recent study found that auditory cortico-striatal synapses were strengthened as rats learned to perform a sound discrimination task (Xiong et al., 2015). Together, these results suggest that posterior striatal circuits undergo major changes in synaptic strength during the initial learning of sound-action association tasks, but do not display such changes when a task requires rapid switching in the associations between sounds and actions without changes in reward (Figure 7). We conclude that after establishing the initial associations, posterior striatal circuits convey sensory information that can be rerouted downstream to drive different responses to the same sensory stimulus.

The conclusion above is consistent with a model in which posterior striatal neurons encode the value of sensory stimuli, so that evoked responses vary significantly when a stimulus becomes predictive of reward (when learning the task for the first time), but do not change when the stimulus predicts the same reward under different stimulus-action contingencies. This is reminiscent of the coding for action-value observed in anterior striatal neurons (Tai et al., 2012). Coding of stimulus-value by posterior striatal neurons would results in a stable representation of sound features when the association between these stimuli and rewards remains constant, even when animals must modify their behavioral responses in order to obtain reward.

## Materials and Methods

### Animal subjects

17 adult male wild-type mice (C57/BL6J) and 7 transgenic adult male mice were used in this study. Transgenic mice expressing Cre recombinase under control of the dopamine D1 receptor (036916-UCD from MMRRC) were crossed with LSL-ChR2 mice (012569 from JAX) to produce mice expressing ChR2 in D1-positive neurons in the striatum (Drd1a::ChR2 mice). Mice had ad libitum access to food, but water was restricted. Free water was provided on days with no experimental sessions. All procedures were carried out in accordance with National Institutes of Health standards and were approved by the University of Oregon Institutional Animal Care and Use Committee.

### Study design and statistics

Sample sizes were based on previous literature in the field (Znamenskiy and Zador, 2013; Jaramillo et al., 2014). Nonparametric statistical tests were used with no assumption of normality of the sample distributions. When comparing laser stimulation in transgenic *vs.* wild-type mice, the experimenter was not blind to the genotype of animals, but data collection was automated to minimize potential biases.

### Behavioral task

The two-alternative choice sound discrimination task was carried out inside single-walled sound-isolation boxes (IAC-Acoustics). Behavioral data was collected using the taskontrol platform (www.github.com/sjara/taskontrol) developed in our laboratory using the Python programming language (<u>www.python.org</u>). Mice initiated each trial by poking their noses into the center port of a three-port behavior chamber. After a silent delay of random duration (150-250 ms, uniformly distributed), a narrow-band sound (chord) was presented for 100 ms. Animals were required to stay in the center port until the end of the sound and then chose one of the two side ports for reward (2 ul of water) according to the frequency of the sound (low-frequency: left port; high-frequency: right port). If animals withdrew before the end of the stimulus, the trial was aborted and ignored in the analysis. Stimuli were chords composed of 12 simultaneous pure tones logarithmically spaced in the range f/1.2 to 1.2f for a given center frequency f. Within a behavioral session, we used 6 or 8 distinct center frequencies. The intensity of all sound components was set to the same value between 30-50 dB-SPL (changing from one trial to the next) during the initial training, but fixed during testing at 50 dB-SPL. Each behavioral session lasted 60 to 90 minutes.

Switching task: To test mice under changing stimulus-action associations, we used a variation of the two-alternative task described above (Jaramillo and Zador, 2014). A single session consisted of several blocks of 200-250 trials, with 2 (out of 3) possible sound stimuli presented in each block. In a ‘low boundary block’, mice were required to choose the left reward port after a low-frequency sound, and choose the right port for a middle-frequency sound. In a ‘high boundary block’, mice were required to discriminate between the middle-frequency sound and a high-frequency sound, with the middle-frequency sound now being rewarded on the left port (Figure 7A). No additional cue was given indicating the change in contingency. The initial contingency in a session was randomized from one day to the next.

### Surgical implant for tetrode arrays

Animals were anesthetized with isoflurane through a nose cone on the stereotaxic apparatus. Mice were surgically implanted with a custom-made microdrive containing eight tetrodes targeting the right posterior striatum. Each tetrode was composed of four tungsten wires (CFW0011845, California Fine Wire) twisted together. The eight tetrodes varied in length with 500 um difference between the longest and the shortest tetrodes. Tetrodes were positioned at 1.7 mm posterior to bregma, 3.5-3.55 mm from midline, and 2 mm from the brain surface at the time of implantation. All animals were monitored and recovered fully before behavioral and electrophysiological experiments.

### Neural recordings

Electrical signals were collected using an RHD2000 acquisition system (Intan Technologies) and OpenEphys software (<u>www.open-ephys.org</u>). Evoked responses to sound were monitored daily and tetrodes were moved down after each recording session. At the first depth where sound-evoked responses were observed, we started collecting electrophysiological data during the sound discrimination task. Recordings for each animal stopped when no more sound responses were observed. Tetrode locations were confirmed histologically based on electrolytic lesions and fluorescent markers. Electrodes were coated with DiI (Thermo Fisher Scientific, Cat#V22885) before implanting.

### Optogenetic stimulation in awake mice

Optical fibers (CFML12U-20, ThorLabs) were cleaved and etched with hydrofluoric acid for 40 minutes to obtain a cone-shaped tip. Each optical fiber was glued to a metal guide tube that helped secure the fiber ferrule to the skull. Before implant, optical fibers were connected to a blue laser (445 nm) built in-house and the light output calibrated using a PM100D power meter (ThorLabs). Drd1a::ChR2 mice were implanted with optical fibers bilaterally targeting either the anterior dorsal striatum (0.5 mm anterior to bregma, 1.6 mm from midline and 2.5 mm from brain surface) or the posterior striatum (1.7 mm posterior to bregma, 3.5 mm from midline and 2.1 mm from brain surface). Optical fibers implanted in the posterior striatum were attached to a movable microdrive together with a tetrode bundle, and were moved down between stimulation sessions. Fiber locations were verified histologically postmortem.

To assess the effect of stimulating striatal direct-pathway neurons on movement initiation, each mouse was placed in a square activity chamber (22 cm by 22 cm) and video-recorded from a camera mounted above the chamber. Two pieces of colored tape were placed on the implant to facilitate tracking. Optical stimulations were carried out unilaterally at 1 mW (measured at the tip of the fiber before implantation) for 1.5 seconds each time. Changes in the angle of the mouse’s head were tracked across video frames (see Analysis of behavioral data for details on this analysis) using custom software written in Python (<u>www.python.org</u>) and OpenCV (opencv.org). To assess the effect of activating posterior striatal direct-pathway neurons during sound discrimination, unilateral optical stimulations were carried out while animals performed the two-alternative choice task. To assess the frequency preference of each stimulation site, neural recordings were made at the depth of the optical fiber via the attached tetrode bundle, while animals were presented with sound stimuli similar to those used in the task before each behavioral session. Each behavioral session lasted for 80-120 minutes during which animals performed 500-900 trials; laser stimulations were delivered in 20% of the trials. In a stimulation trial, the laser stimulation (1 mW, 200 ms) started 50 ms before the sound onset and ended 50 ms after the sound.

### Muscimol inactivation

Bilateral craniotomies were performed under stereotactic surgery over the posterior striatum (1.7 mm posterior to bregma, 3.55 mm lateral from midline) of mice trained in the two-alternative choice sound discrimination task. Headbars were implanted to allow for head-fixation. Each craniotomy was protected with a plastic ring and filled with silicon elastomer (Sylgard 170, Dow Corning). Animals were allowed to recover for at least 3 days before resuming behavioral training. Following recovery, implanted animals were trained on the sound discrimination task until they reached their pre-surgery performance level before beginning muscimol inactivation.

For intracranial injection, we used glass pipettes (5 ul Disposable Micropipettes, VWR) pulled and trimmed to an inner diameter of 15-20 micrometers at the tip. Animals were head-fixed and allowed to run on a wheel during the injection. Craniotomies were exposed by removing the silicon elastomer covering, and a glass pipette filled with reagent (either muscimol or saline) was lowered into the brain to a depth of 3.1mm from brain surface using a micromanipulator. A volume of 45 nl of muscimol (0.25 mg/ml, final dose of 11.25 ng/hemisphere) was injected under air pressure in each hemisphere at a rate of 90 nl/min. The pipette was left in place for 60 seconds following the injection, then raised 0.5mm and left in place for another 60 seconds before being removed. Injection in the second hemisphere was always completed within 10 minutes of the first injection. The craniotomies were then protected with a new silicon elastomer cap, and the mouse was placed back into its home cage for 30 minutes before starting the behavior session. After collection of 4 saline sessions and 4 muscimol sessions, 45 nl of fluorescent dye (DiI, Thermo Fisher Scientific) was injected at the same injection coordinates. Animals were then perfused transcardially with 5% paraformaldehyde, and brains were extracted and postfixed for 12-24 hours. Brains were then sliced (100 um) and imaged to verify the location of fluorescent dye injection.

Fluorescent muscimol (Muscimol, BODIPY TMR-X Conjugate, Thermo Fisher Scientific) was dissolved in phosphate-buffered saline to a final concentration of 0.5 mg/ml. We followed the same protocol for intracranial injection as for muscimol. The injection volume was 360 nl/hemisphere (final dose of 180 ng/hemisphere) to account for the larger molecular weight and reduced spread of fluorescent muscimol. As with muscimol injections, animals rested for 30 minutes before starting the behavior session. Animals were euthanized and transcardially perfused with 5% paraformaldehyde within 1 hour of finishing behavioral testing. Brains were extracted and postfixed overnight, and sliced (100 um) to quantify spread of fluorescent muscimol.

### Analysis of behavioral data

Estimation on head angle: Centroids for the green and red colored tape attached to the left and right side of each animal’s implant were estimated for each video frame. Head angle (Figure 1B) was calculated from the line connecting these centroids. Head angle for frames in which colored tape was not visible was estimated using linear interpolation from other frames. Total change in head angle for each trial (in Figure 1C) was calculated as the angle at the end of the stimulation minus the angle at the beginning of the stimulation.

Psychometric curve fitting for Figure 2 and Figure 3 was performed via constrained maximum likelihood to estimate the parameters of a logistic sigmoid function (<u>http://psignifit.sourceforge.net</u>).

### Analysis of neuronal data

Data were analyzed using in-house software developed in Python (<u>www.python.org</u>). Spiking activity of single units was isolated using an automated expectation maximization algorithm (Klustakwik; Kadir et al., 2013). Isolated clusters were only included in the analysis if less than 2% of inter-spike intervals were shorter than 2 ms. Spike shapes were manually inspected to exclude noise signals.

A cell was considered sound-responsive if activity evoked by all sound frequencies pooled together (0 to 100 ms from sound onset) was significantly different from baseline spontaneous activity (-100 to 0 ms from sound onset) according to a Wilcoxon rank-sum test. The sound response index for each cell was calculated as (S-B)/(S+B), where S is the average sound-evoked response and B is the average spontaneous firing rate. To determine whether neuronal firing encoded task-specific sound features, we calculated a selectivity index which quantified the difference between the firing rates in response to high-versus low-frequency sounds with respect to the categorization boundary: (*F*_*high*_ *- F*_*low*_)*/*(*F*_*high*_ + *F*_*low*_).

For movement-related responses, average firing rates in a 50-150 ms window after the animal exited the center port were quantified. A movement modulation index was calculated by (I - C)/(I + C), where I and C were average firing rates from trials with movement to the reward port ipsilateral and contralateral to the recording site, respectively. To ask whether neuronal activity encoded movement direction, firing rates for trials with contralateral versus ipsilateral movement were compared using Wilcoxon rank-sum test; neurons with a resulting p value of less than 0.05 were considered ‘movement direction selective’.

To evaluate whether neuronal response predicted the animal’s choice, we focused on neurons that showed sound-evoked response to the stimuli with ambiguous behavioral choices (stimuli near discrimination boundary in the sound discrimination task) or periodically updated categorization contingencies (intermediate frequency stimulus in the switching task). To assess responsiveness to the stimuli of interest, spike counts were quantified in non-overlapping bins of 25 ms during the response period (0-100 ms after sound onset) and during the baseline period (25-50 ms before sound onset). A test statistic (z-score) was computed for each response bin in relation to the baseline bin using a Wilcoxon rank-sum test. We considered a cell responsive if the z-score of any bin during the response period fell outside the range (-3,3). To quantify the impact of choice on neuronal activity, we calculated a modulation index (MI) using the equation: (I - C)/(I + C), where I and C were average firing rate in trials with ipsilateral and contralateral choices, respectively. Only sessions with at least 60% correct trials were included in the switching task analysis. To ensure that the MI reflected neuronal activity during stable performance of the task, the first 20 trials after a contingency switch were excluded and only correct trials were included in the MI calculation. We tested statistical significance of choice modulation for each cell using the Wilcoxon rank-sum test between the evoked firing of each choice (sound discrimination task) or contingency (switching task). In the sound discrimination task, this analysis focused on the sound stimulus that elicited the largest sound-evoked response out of the two most ambiguous stimuli closest to the categorization boundary. To exclude effects of nonstationarity fluctuations in neuronal firing rate over time in the switching task, cells were counted as significantly modulated only if the modulation effect was observed in at least two different switches of contingency blocks.

## Author contributions

S.J. conceived the project. S.J., W.I.W., L.G., and N.D.P. designed the experiments. W.I.W., P.L.P., and L.G. conducted and analyzed the electrophysiological recordings. L.G. conducted and analyzed the optogenetic studies. N.D.P. conducted and analyzed the muscimol inactivation studies. S.J. supervised all aspects of the work. L.G., N.D.P., and S.J. wrote the paper.

## Acknowledgement

The authors would like to thank members of the Jaramillo lab for discussion and comments on the manuscript. This research was supported by the National Institute on Deafness and Other Communication Disorders (R01 DC015531), the Medical Research Foundation, and the Office of the Vice President for Research & Innovation at the University of Oregon.

---

**Figure S1.**
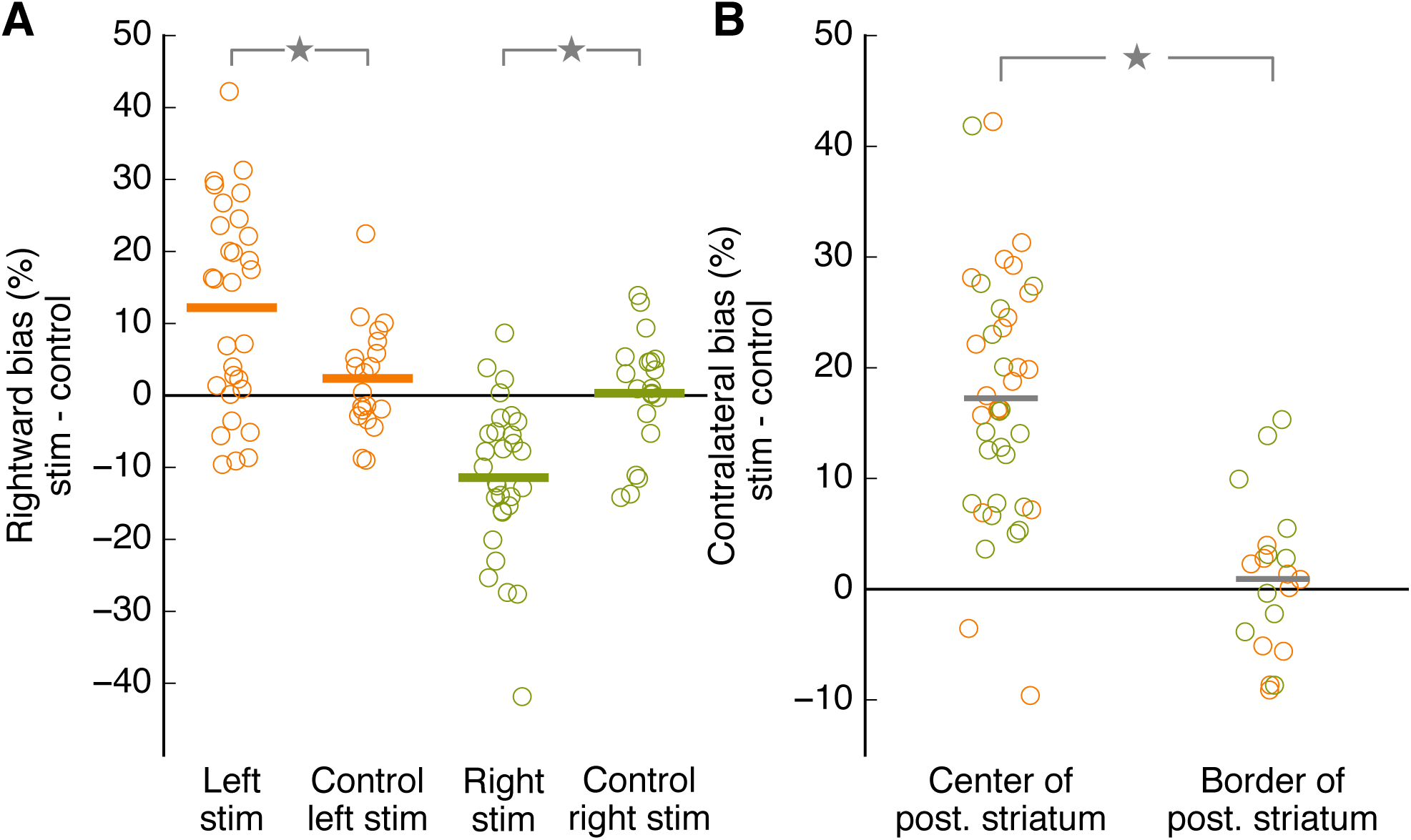
Laser stimulation did not produce behavioral bias in wild-type mice. Related to Figure 2. (A) Change in the percentage of rightward choices during laser stimulation for each hemisphere in Drd1a::ChR2 mice and wild-type control mice. Each dot represents one session (Drd1a::ChR2: N=3 mice, 10 sessions each hemisphere per mouse; wild-type control: N=2 mice, 10 sessions each hemisphere per mouse). Horizontal bars represent averages across all sessions for all mice. Stimulation produced significantly different biases in the Drd1a::ChR2 mice versus the control mice in each hemisphere (*p* = 0.028 left, *p <* 0.001 right, Wilcoxon rank-sum test). (B) Comparison of contralateral bias resulting from activation of different sites on the medial-lateral axis of posterior striatum in Drd1a::ChR2 mice. Stimulation of sites at the center of the striatum produced significantly higher bias than sites near the border between striatum and cortex (*p <* 0.001, Wilcoxon rank-sum test).

**Figure S2.**
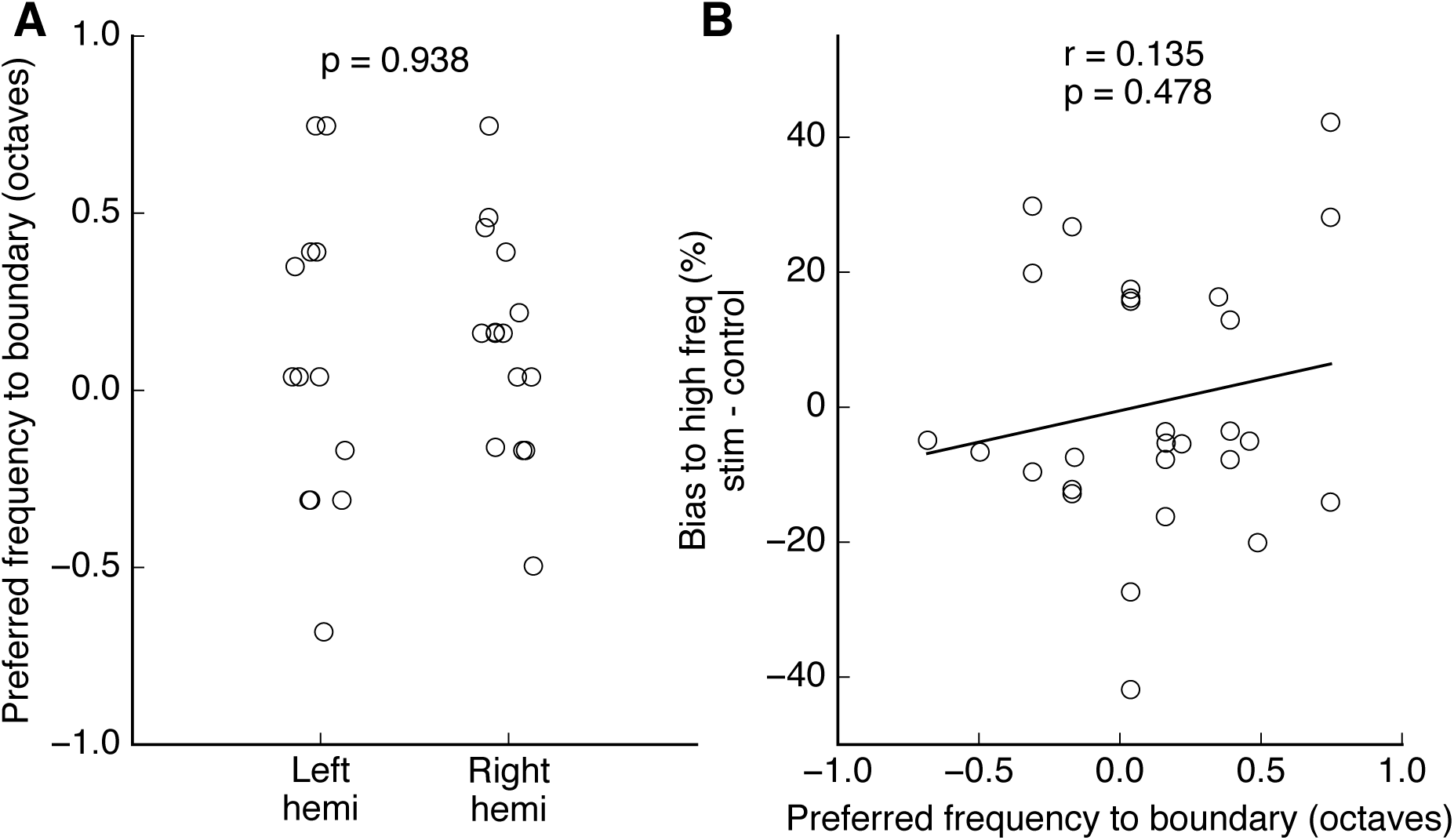
Behavioral bias did not correlate with frequency preference of optical stimulation sites. Related to Figure 2. (A) Preferred sound stimuli (relative to categorization boundary) at optogenetic stimulation sites in each hemisphere of Drd1a::ChR2 mice. Each dot represents a stimulus selective site. Frequency tuning was not significantly different between hemispheres (*p* = 0.938, Wilcoxon rank-sum test). (B) There was no significant correlation between choice bias and frequency tuning at a stimulated site (*r* = 0.135, *p* = 0.478, Spearman correlation test).

**Figure S3.**
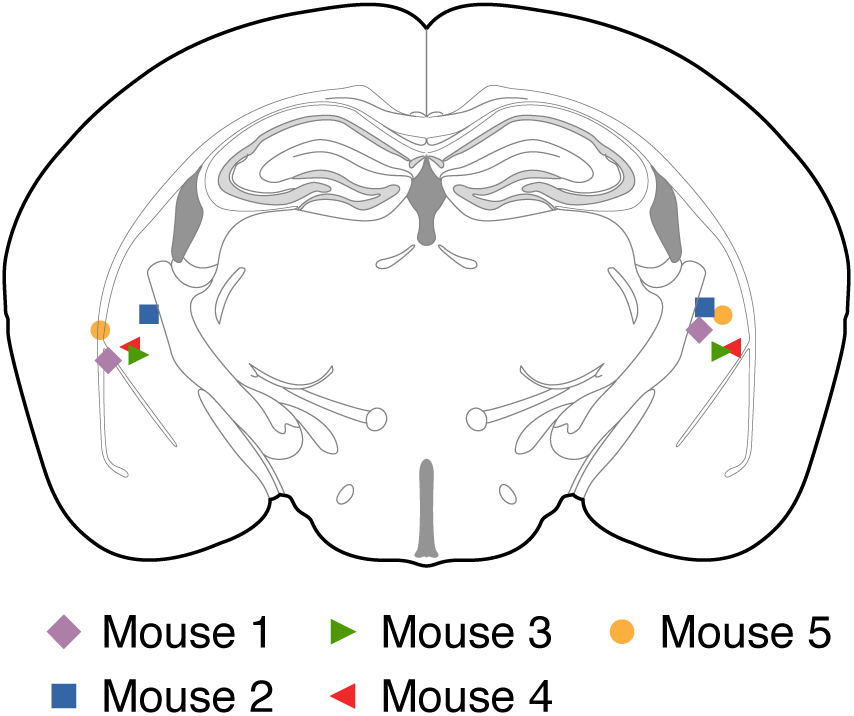
Center of muscimol injection in the posterior striatum for each mouse tested. Related to Figure 3. After the last behavioral session, a fluorescent dye was injected and the center of the injection estimated from fixed brain sections.

**Figure S4.**
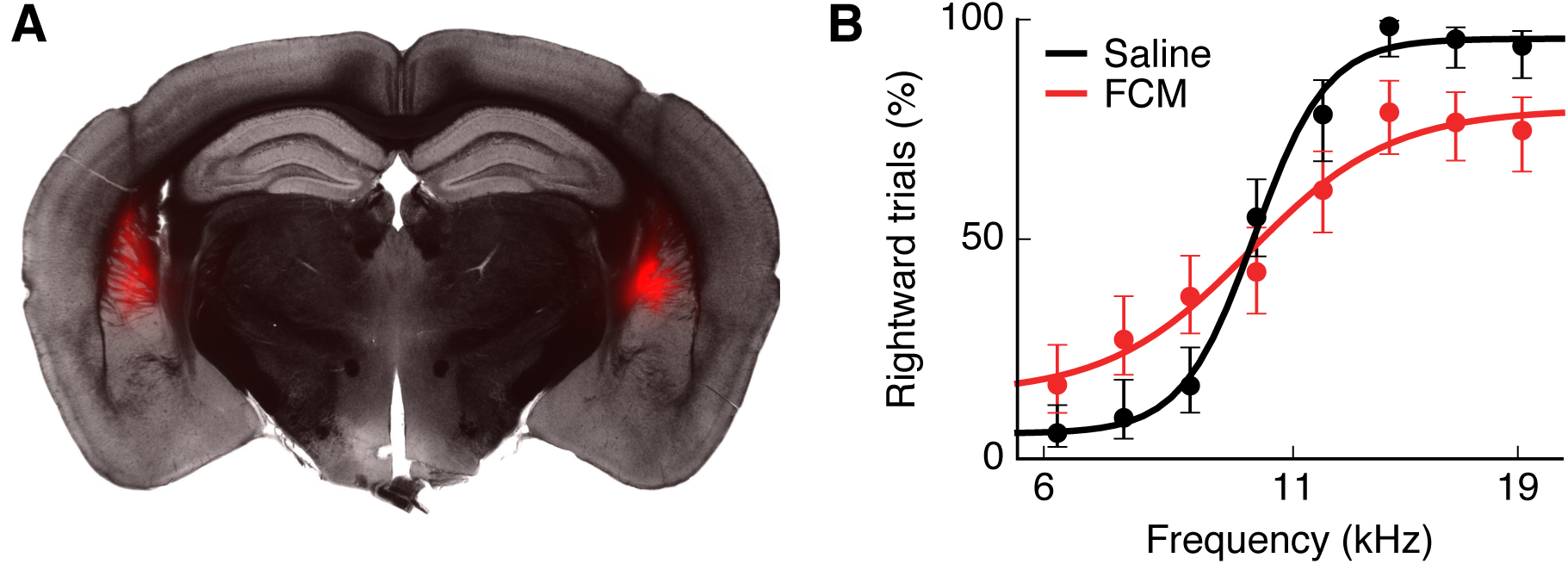
Inactivation of posterior striatum using fluorescent muscimol impaired task performance. Related to Figure 3. (A) Coronal section showing injection site and spread of Muscimol-BODIPY TMR-X Conjugate (Fluorescent Conjugated Muscimol, FCM) in the posterior striatum. (B) Average psychometric performance of the mouse in (A) after injection of FCM (red, 1 session) or saline control (black, 1 session). Error bars indicate 95% confidence intervals.

**Figure S5.**
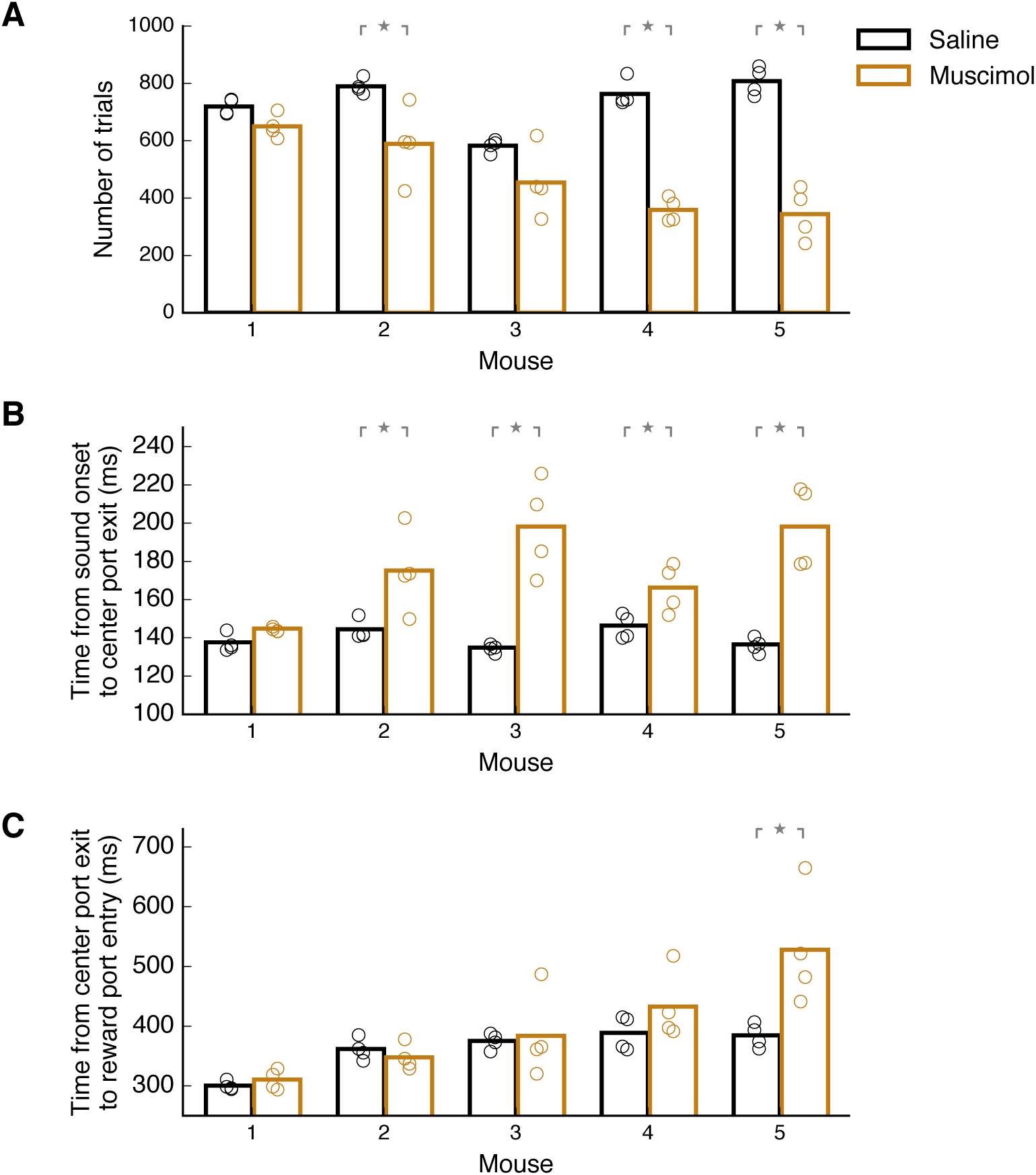
Effect of muscimol inactivation of posterior striatum on movement during the sound discrimination task. Related to Figure 3. (A) Number of trials performed in 1 hour on each saline session (black) and each muscimol session (brown) for each mouse. Bars indicate average across sessions for each mouse. Muscimol inactivation significantly reduced the number of trials performed by 3 out of 5 mice (*p* = 0.021, Wilcoxon rank-sum test), but did not preclude animals from performing hundreds of trials in each session. (B) Average withdrawal time from center port after sound onset on each session. Muscimol inactivation led to slower withdrawals from the center port for four out of five mice (*p <* 0.05, Wilcoxon rank-sum test) compared to saline sessions. (C) Average movement time from center port to side reward ports on each session. A change in movement speed was observed in only one out of five mice (*p <* 0.05, Wilcoxon rank-sum test).

**Figure S6.**
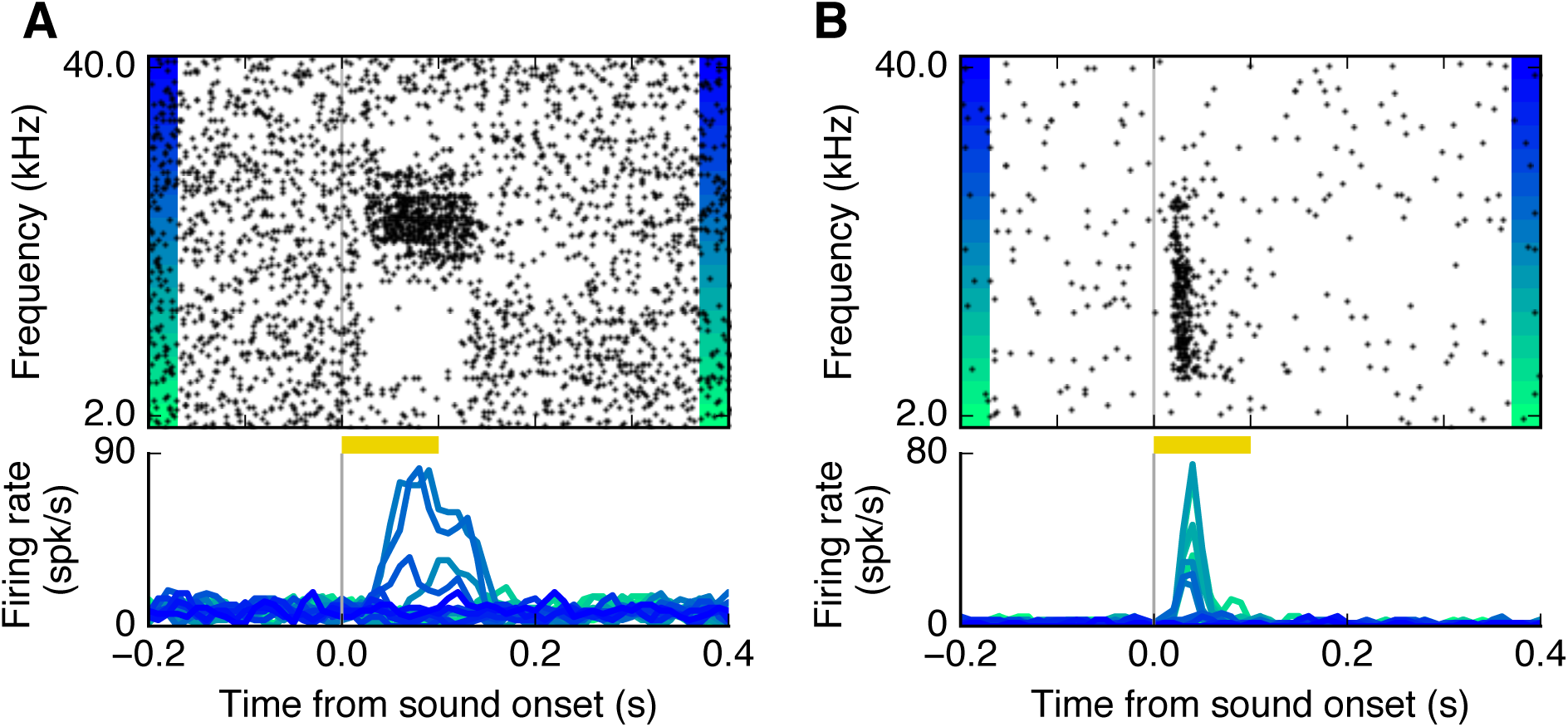
Posterior striatal neurons displayed frequency-selective sound-evoked responses outside the sound discrimination task. Related to Figure 4. (A,B) Examples of sound responses from two different posterior striatal neurons to different frequencies of sound outside the sound discrimination task. Yellow bar indicates the duration of the sound. Cells showed clear frequency selectivity.

**Figure S7.**
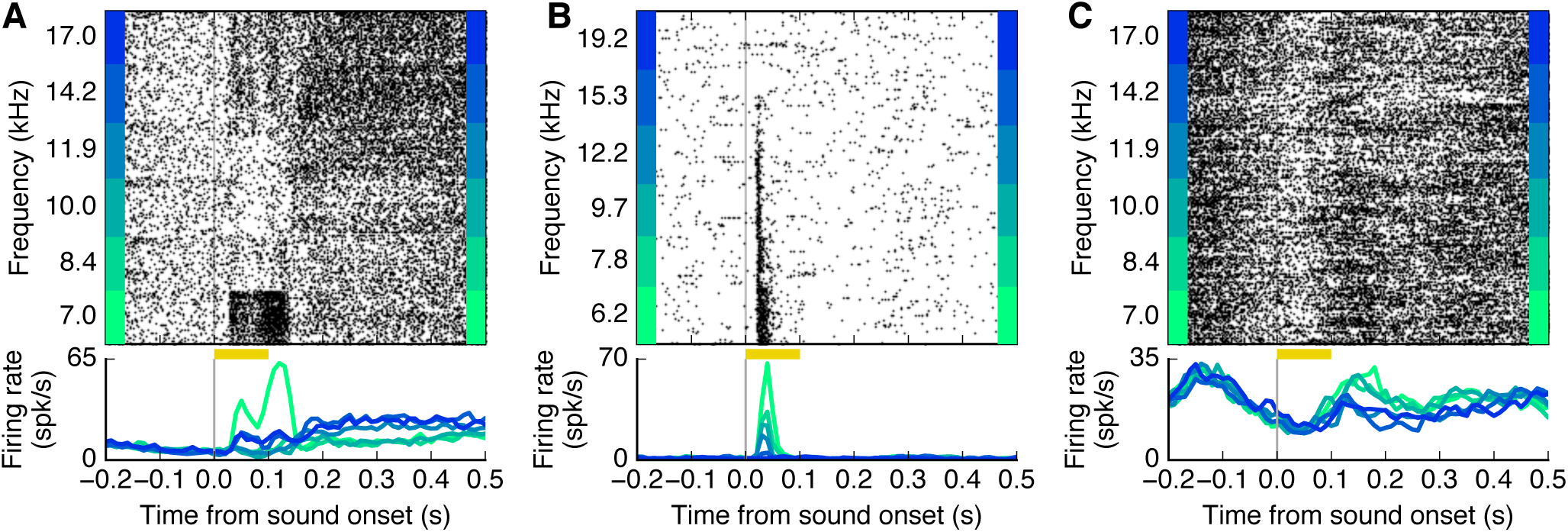
Neuronal encoding of sound stimuli and movement direction in the posterior striatum. Related to Figure 4 and Figure 5. (A) Example cell that was both selective to sound frequency and movement direction. The plot includes trials in which the mouse went to the left port for the 3 lower frequencies and to the right port for the 3 higher frequencies. (B) Example cell that responded to sound stimuli, but was not sensitive to movement direction. (C) Example cell that did not respond to sound but was selectively active when animal was moving to one side port versus the other.

**Figure S8.**
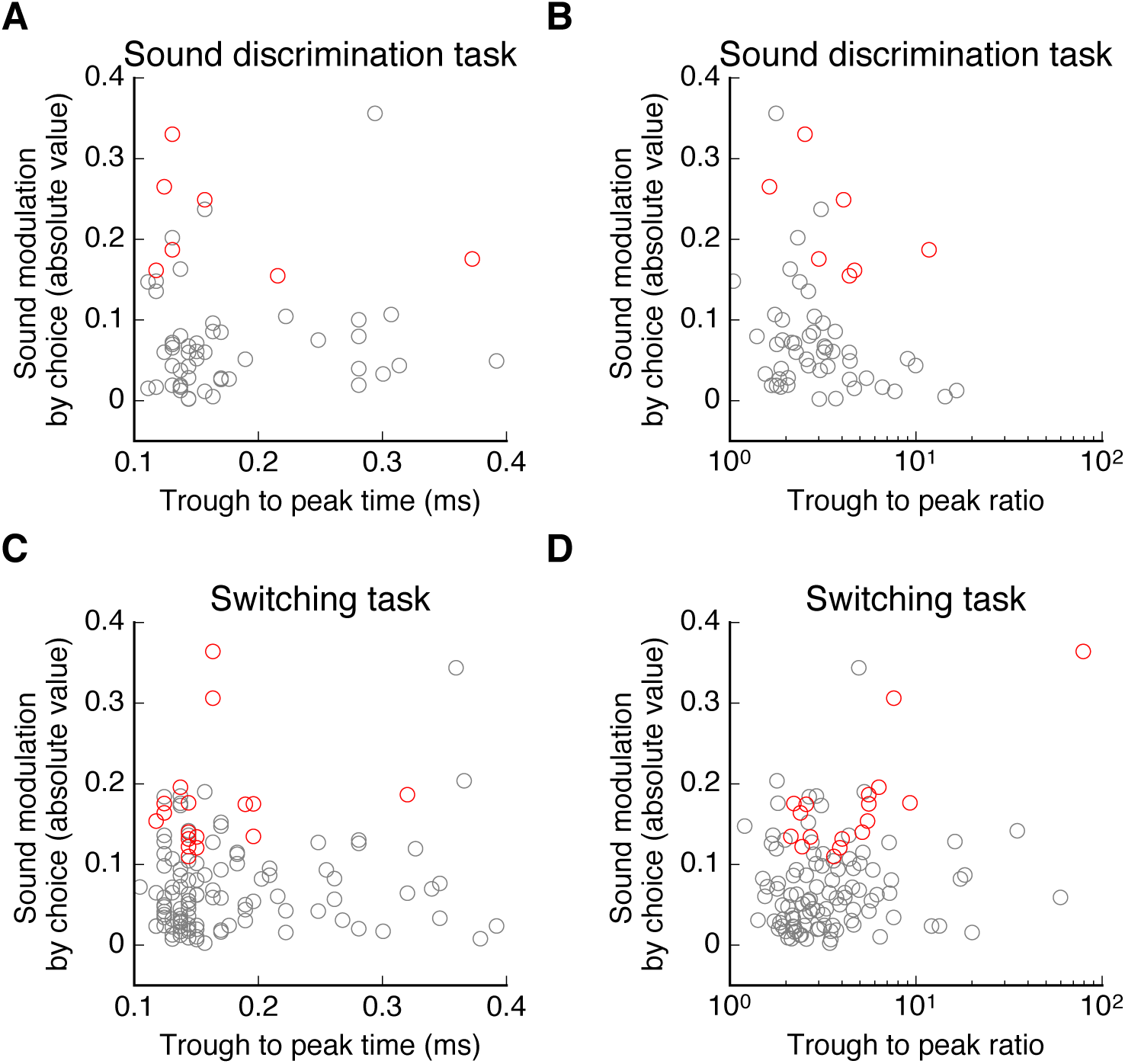
Spike waveform parameters did not predict whether a cell’s sound response was modulated by choice. Related to Figure 6 and Figure 7. (A) Spike trough-to-peak time was not correlated to the modulation index of sound by choice in the case of ambiguous stimuli (N = 67 cells from 5 mice, *r* = −0.02, *p* = 0.8, Spearman correlation test). (B) Modulation index of a cell was not correlated with its trough-to-peak ratio (N = 67 cells from 5 mice, *r* = −0.2, *p* = 0.09, Spearman correlation test). (C-D) Similarly, when mice were required to rapidly update stimulus-action association (switching task), sound response modulation index was not correlated with either trough-to-peak time (*r* = 0.06, *p* = 0.5, Spearman correlation test) or trough-to-peak ratio (N = 155 cells from 4 mice, *r* = 0.17, *p* = 0.06, Spearman correlation test).

**Figure S9.**
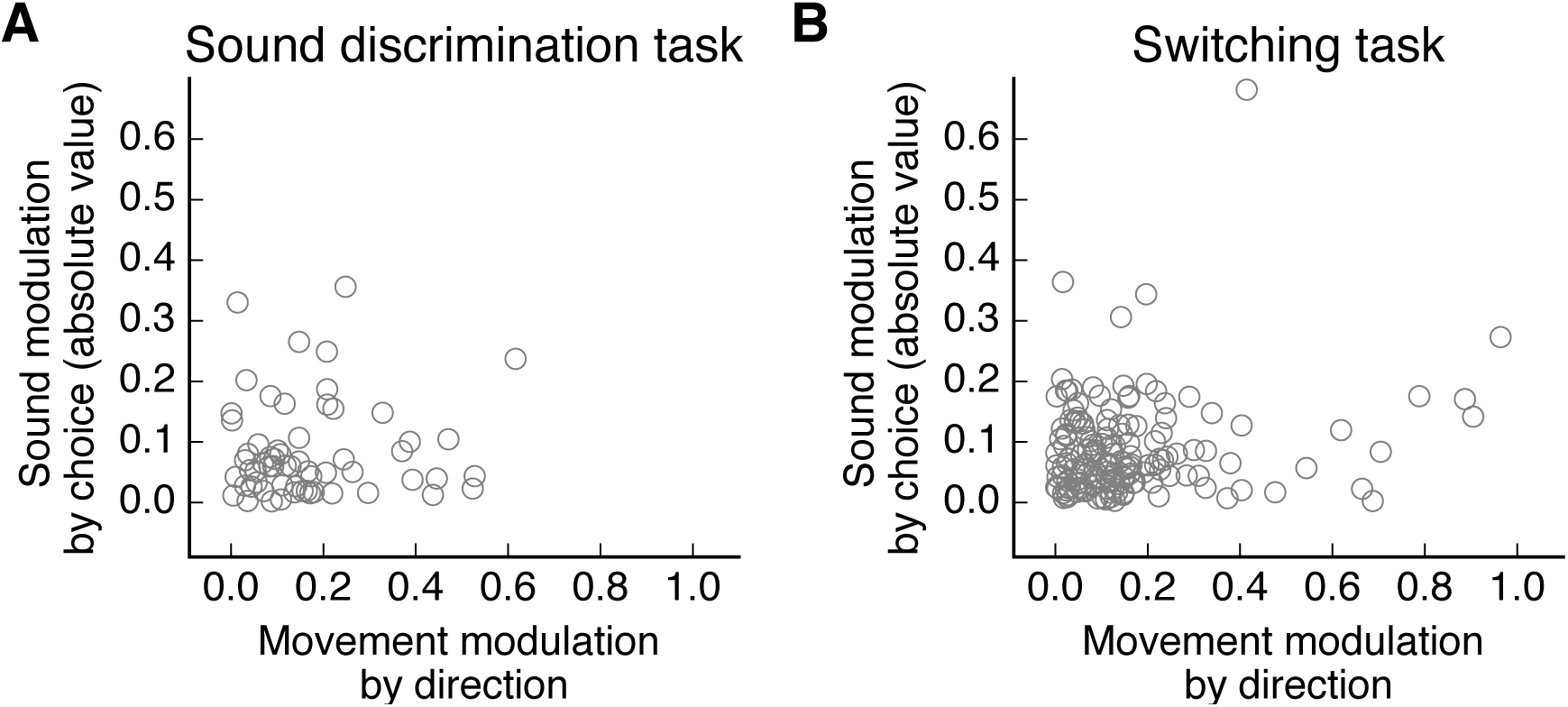
Movement direction selectivity did not predict whether a cell’s sound response was modulated by choice. Related to Figure 6 and Figure 7. (A) For cells that were responsive to sound stimuli near the categorization boundary, the magnitude of movement selectivity index did not correlate with that of the sound modulation index (N = 67 cells from 5 mice, *r* = 0.02, *p* = 0.8, Spearman correlation test). (B) For cells responsive to the stimulus that reversed associated in the switching task, the magnitude of movement selectivity index did not correlate with that of the sound modulation index (N = 155 cells from 4 mice, *r* = 0.07, *p* = 0.4, Spearman correlation test).

**Figure S10.**
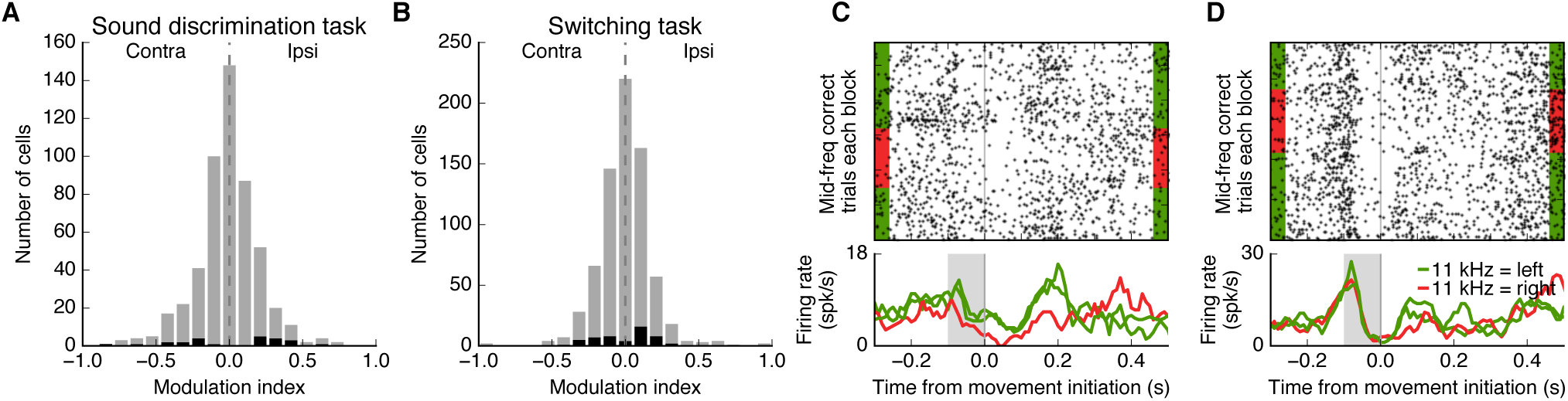
Neuronal activity prior to movement initiation did not predict choice. Related to Figure 6 and Figure 7. (A) Influence of choice on neuronal activity before action initiation for ambiguous sound stimulus in the sound discrimination task (N = 520 cells from 5 mice). Less than 5% of neurons showed a significant change in activity before movement predictive of subsequent choice (*p <* 0.05, Wilcoxon rank-sum test), shown in black. (B) Influence of changing sound-action association for the stimulus of intermediate frequency on neuronal activity before action initiation (N = 725 cells from 4 mice). Only 7% of neurons showed a significant change across sound-action contingencies (*p <* 0.05, Wilcoxon rank-sum test), shown in black. (C) Responses of one posterior striatal neuron aligned to movement start (0 on x-axis) in trials with the stimulus of intermediate frequency for three blocks of trials (switching task). Activity in the 0-100 msec window before movement initiation (shaded area) for this neuron changed systematically depending on the rewarded action associated with the stimulus. (D) Pre-movement activity of a different neuron showing no change across sound-action contingencies.

